# A repository of Ogden syndrome patient derived iPSC lines and isogenic pairs by X-chromosome screening and genome-editing

**DOI:** 10.1101/2024.09.28.615067

**Authors:** Josephine Wesely, Tom Rusielewicz, Yu-Ren Chen, Brigham Hartley, Dayna McKenzie, Matthew K. Yim, Colin Maguire, Ryan Bia, Sarah Franklin, Rikhil Makwana, Elaine Marchi, Manali Nikte, Soha Patil, Maria Sapar, Dorota Moroziewicz, NYSCF Global Stem Cell Array® Team, Lauren Bauer, Jeannie T. Lee, Frederick J. Monsma, Daniel Paull, Gholson J. Lyon

## Abstract

Amino-terminal (Nt-) acetylation (NTA) is a common protein modification, affecting 80% of cytosolic proteins in humans. The human essential gene, *NAA10,* encodes the enzyme NAA10, as the catalytic subunit for the N-terminal acetyltransferase A (NatA) complex, including the accessory protein, NAA15. The first human disease directly involving *NAA10* was discovered in 2011, and it was named Ogden syndrome (OS), after the location of the first affected family residing in Ogden, Utah, USA. Since that time, other variants have been found in *NAA10* and *NAA15.* Here we describe the generation of 31 iPSC lines, with 16 from females and 15 from males. This cohort includes CRISPR-mediated correction to the wild-type genotype in 4 male lines, along with editing one female line to generate homozygous wild-type or mutant clones. Following the monoclonalizaiton and screening for X-chromosome activation status in female lines, 3 additional pairs of female lines, in which either the wild type allele is on the active X chromosome (Xa) or the pathogenic variant allele is on Xa, have been generated. Subsets of this cohort have been successfully used to make cardiomyocytes and neural progenitor cells (NPCs). These cell lines are made available to the community via the NYSCF Repository.

## 1. Introduction

Amino-terminal (Nt-) acetylation (NTA) is a form of co-translational or post-translation modification that has been conserved across multiple eukaryotic species and is present across 80% of all proteins in humans^1^. Functionally, NTA works by irreversibly incorporating an acetyl group to the Nt residue of a protein. There are currently eight N-terminal acetyltransferases (NATs) in eukaryotes named NatA-H. Each NAT is composed of a catalytic domain and in many cases a single or several auxiliary domains that serve various functions ranging from ribosome binding to substrate-specificity modification^2^. Most NAT catalytic subunits display substrate specificity based on the first few residues after the N-terminal residue^3^. NTA functions to alter protein complex interactions^4^, trafficking^5^, folding^6^, and degradation^7^.

NatA is thought to acetylate around 40% of the proteome in humans^1^. NatA is a heterodimer composed of the NAA10 catalytic subunit and the NAA15 auxiliary subunit^8^ that interacts with several chaperone proteins including NAA50/NatE and HYPK^9–11^. NAA15 facilitates these interactions as well as anchors NAA10 to the ribosome for function, with HYPK serving to inhibit the NatA complex activity until necessary^11–13^. The NatA complex acetylates the following second residues of the nascent protein chain, serine, glycine, alanine, threonine and cysteine residues, after methionine removal^14^.

Pathogenic variants of both NAA10 and NAA15 are associated with several pathological phenotypes. Pathogenic variants of NAA10 have been implicated in various disease states including cancer^15–23^, Parkinson disease^24,25^, and NAA10 related neurodevelopmental syndrome, colloquially known as Ogden Syndrome (OS). OS is an X-linked neurodevelopmental syndrome first characterized in 2011 in a family in Ogden, Utah associated with a p.Ser37Pro missense pathogenic variant that manifested as developmental delay, cardiac abnormalities, distinct facial atypia, and hypotonia^26,27^. Since then, the number of pathogenic variants associated with the disease have increased with there being over 100 confirmed cases observed globally^28,29^. Additional phenotypic manifestations have been observed including sensory abnormalities, gastrointestinal abnormalities, skeletal malformations, and disruptions of the metabolic system with more severe presentations appearing in males than females.^8,28–45^. NAA15-related neurodevelopmental disorder has been shown to usually be milder than OS, where it is characterized by variable penetrance of developmental delay, cardiac abnormalities, and various motor and other functional delays^28,30,46–55^.

Induced pluripotent stem cells (iPSCs) are somatic cells that have been reprogrammed back to an earlier stage of development to allow for subsequent differentiation into different cell types to allow for the study of disease *in vitro*^56^. These iPSCs can be made from somatic cells, such as skin-derived fibroblasts or blood, taken directly from patients with any particular disease, thus allowing the capture and modeling of that particular genetic background, including any pathogenic variants predisposing to disease. Previous iPSC studies derived from Ogden Syndrome patient cells allowed for the generation of cardiomyocytes that, in conjunction with electrophysiological techniques, allowed for the characterization of the long QT phenotype that presents itself in those individuals^33^. The purpose of this paper is to report the creation of a repository of iPSCs from a diverse cohort of patients with *NAA10*-related or *NAA15*-related neurodevelopmental syndrome, representing different pathogenic variants associated with the disease. It is hoped that this repository of such cells, open to anyone to request and utilize, will catalyze future experimentation to better understand the function of *NAA10* and *NAA15* in the context of these disease presentations.

## 2. Results

### 2.1. Resource utility and iPSC lines

All human iPSC lines generated (reprogrammed, X-chromosome screened and gene-edited at the NYSCF Research Institute repository) can be accessed through the NYSCF Repository and used to investigate pathological cellular phenotypes associated with pathogenic variants in *NAA10* in patients with Ogden syndrome. An overview of the iPSC generation pipeline is shown in **Figure 1**. A current list of all available iPSC lines (controls, *NAA10*, and *NAA15*) can be found in **Supplementary Table 1**.

**Figure 1.**
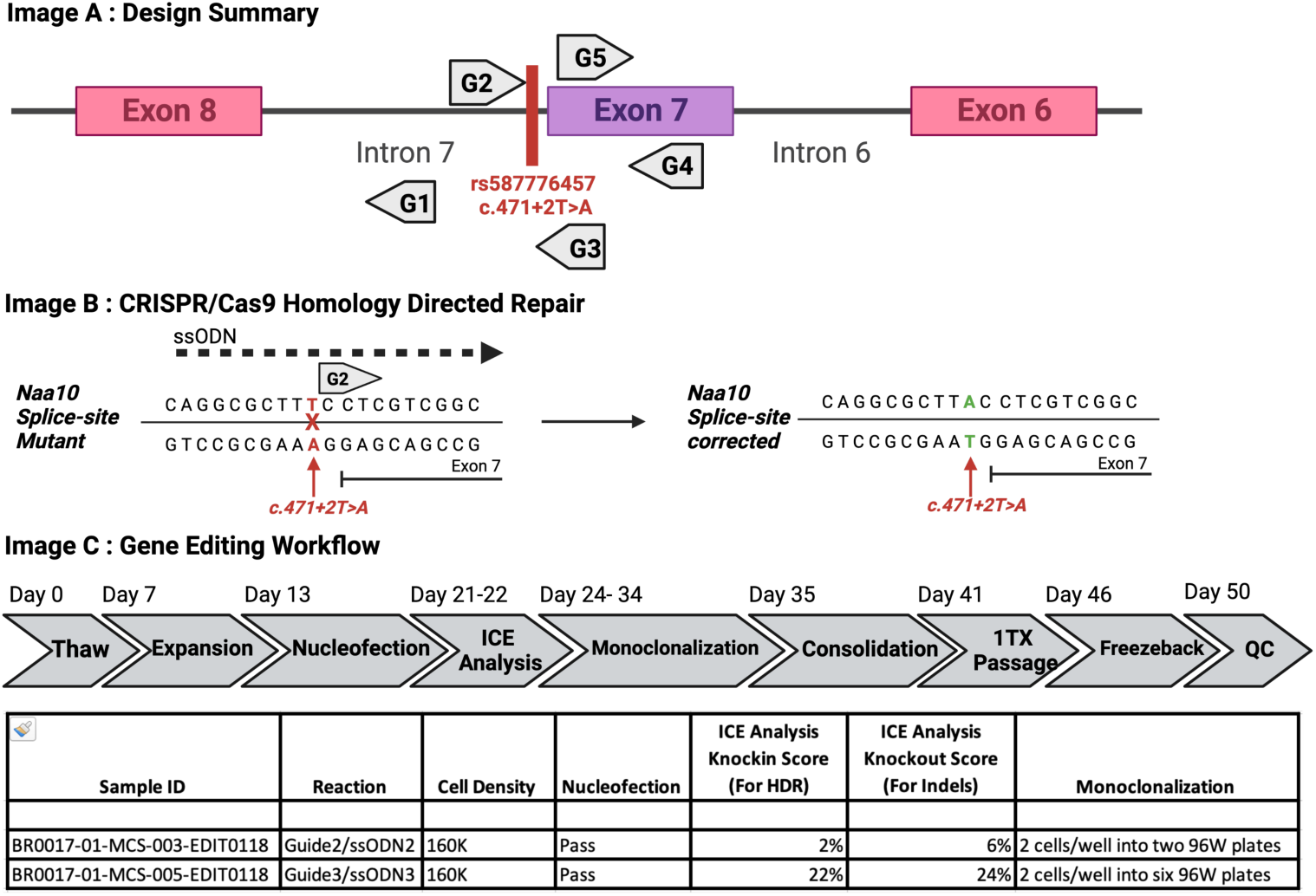
Methodology for the generation of Ogden Syndrome patient derived iPSCs.

### 2.2. X-chromosome screened iPSC lines

*NAA10* is located on the X-chromosome, therefore the status of X-chromosome inactivation in female primary patient samples as well as on the generated iPSC lines was determined for some of the lines. 10 female samples were analyzed of which 5 contained the heterozygous NAA10 Arg83Cys, 3 contained the heterozygous Phe128Leu, 1 contained the heterozygous Ala87Ser and 1 contained the heterozygous Leu121Val. All primary samples were peripheral blood mononuclear cells (PBMCs) except BR0016 which was a fibroblast sample. 7 out of 10 samples showed higher skewing towards the wild type (WT) X-chromosome with a range of 60%-89.9% WT allele expressed, 1 sample showed no skewing (50.5% WT/49.5% pathogenic variant and 2 samples showed skewing towards the pathogenic variant X-chromosome 72.7% and 84% (**Fig. 2A**). Upon reprogramming into iPSC pools, 4 samples did not show a change of skewing (defined by more than 10%), 4 samples shifted more than 10% towards pathogenic variant X-chromosome activation and 2 samples shifted more than 10% towards WT X-chromosome activation (**Fig. 2B** and **Table 1**). We further monoclonalized the iPSC pools and screened all generated monoclonal clones for their X-chromosome activation status. We were able to identify WT X-chromosome activated as well as pathogenic variant X-chromosome activated matching pairs for 3 original patient samples (BR0011, BR0014, BR0016). For 4 samples we only identified WT X-chromosome activated clones (BR0002, BR0004, BR0013, BR0015). For BR0003 we only observed clones with pathogenic variant X-chromosome activation, and for BR0005 we observed mixed activation clones and pathogenic variant X-chromosome activated clones, for BR0012 we observed mixed activation as well as clones with WT X-chromosome activation status **(Table 1).**

**Figure 2.**
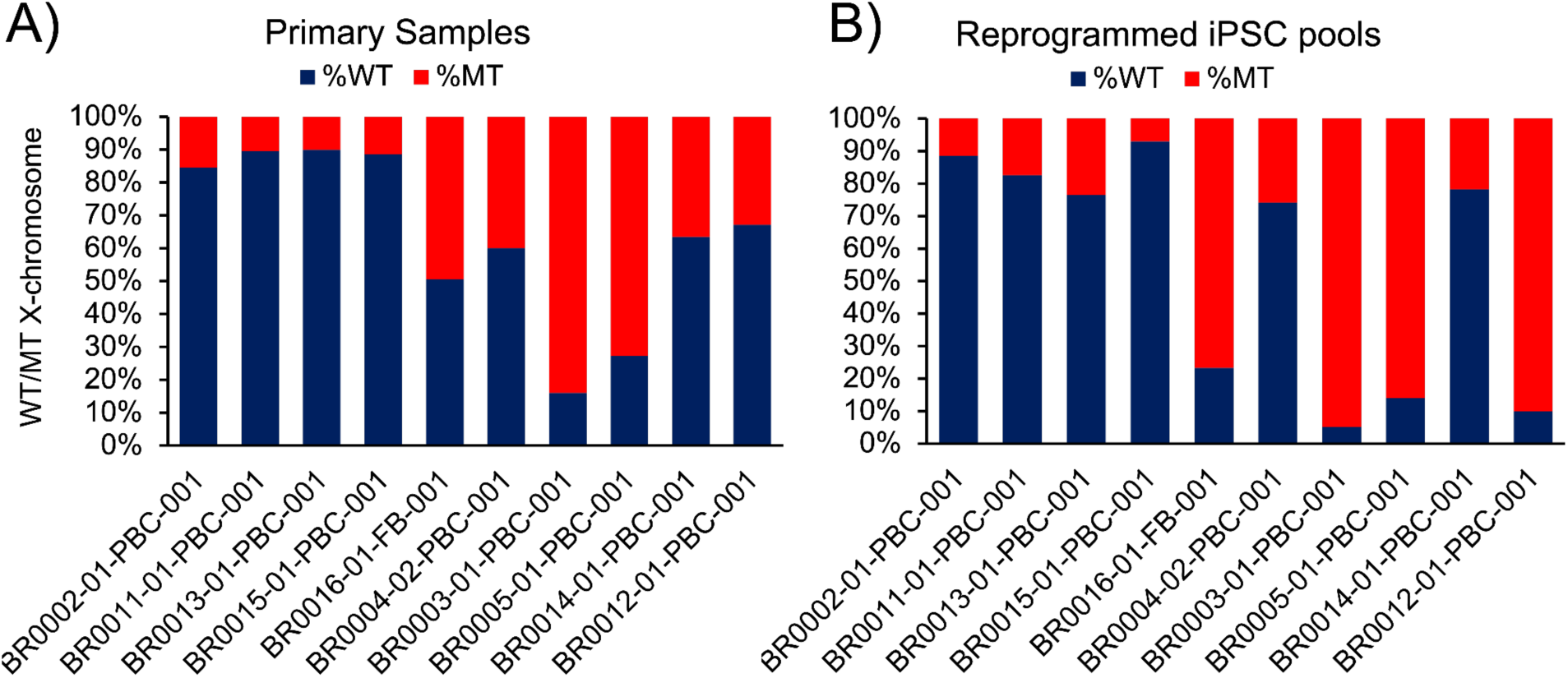
Next Generation Sequencing (NGS) analysis of RNA reverse transcribed into cDNA. A) primary patient samples (PBC indicates PBMCs as source; FB indicates fibroblasts as source) and B) their reprogrammed iPSC pools (%WT=wildtype in blue, %PATHOGENIC VARIANT= mutant in red)

**Table 1.** Percentage WT/pathogenic variant X-chromosome activation in screened iPSC lines.

| NYSCF ID |  | Mutation | Primary samples |  | iPSC pool |  | shift during reprogramming<br>(>10% increase) | iPSC monoclonal lines |  |  |  |
| --- | --- | --- | --- | --- | --- | --- | --- | --- | --- | --- | --- |
|  |  |  | %WT | %MT | %WT | %MT |  | # of lines | # WT | #MT | #WT/MT |
| BR0002-01-PBC-001 | BR0002 | heterozygous p.Arg83Cys | 84.5 | 15.5 | 88.5 | 11.5 | No | 6 | 4 | 0 | 0 |
| BR0011-01-PBC-001 | BR0011 | heterozygous Arg83Cys | 89.5 | 10.5 | 82.6 | 17.4 | No | 8 | 7 | 1 | 0 |
| BR0013-01-PBC-001 | BR0013 | heterozygous Arg83Cys | 89.9 | 10.1 | 76.6 | 23.4 | towards MT | 13 | 13 | 0 | 0 |
| BR0015-01-PBC-001 | BR0015 | heterozygous Arg83Cys | 88.6 | 11.4 | 93 | 7 | No | 3 | 3 | 0 | 0 |
| BR0016-01-FB-001 | BR0016 | heterozygous Arg83Cys | 50.5 | 49.5 | 23.3 | 76.7 | towards MT | 20 | 11 | 9 | 0 |
| BR0004-02-PBC-001 | BR0004 | heterozygous Ala87Ser | 60 | 40 | 74.2 | 25.8 | towards WT | 9 | 8 | 0 | 0 |
| BR0003-01-PBC-001 | BR0003 | heterozygous Phe128Leu | 16 | 84 | 5.1 | 94.9 | No | 8 | 0 | 8 | 0 |
| BR0005-01-PBC-001 | BR0005 | heterozygous Phe128Leu | 27.3 | 72.7 | 14.1 | 85.9 | towards MT | 48 | 0 | 46 | 2 |
| BR0014-01-PBC-001 | BR0014 | heterozygous Phe128Leu | 63.5 | 36.5 | 78.3 | 21.7 | towards WT | 12 | 3 | 9 | 0 |
| BR0012-01-PBC-001 | BR0012 | heterozygous Leu121Val | 67.1 | 32.9 | 10 | 90 | towards MT | 8 | 5 | 0 | 3 |

### 2.3. Genome edited iPSC lines

We generated monoclonal isogenic NAA10-corrected iPSC lines for NAA10 R83C (one male line and one female line), a NAA10 R83C homozygous mutated clone for the female line and c.471+2T→A (splicing site of intron 7, removes exon 7) using asymmetric single-stranded oligo DNA nucleotides (ssODNs) with Cas9 protein/sgRNA ribonucleoprotein complex (Cas9-RNP). The correction for these isogenic lines was confirmed by Sanger-Sequencing. The lines were characterized and validated as described in **Table 2**. Two other male iPSC lines were previously corrected at Stanford, as previously reported^33^, and these lines are now available as part of this cohort, in the NYSCF Repository. The single nucleotide variant (SNV) corrected lines showed a typical iPSC morphology and were karyotypically identical to the parental line suggesting morphological equivalency. Pluripotency was evaluated by gene expression analysis of the pluripotency markers including NANOG, SOX2, POU5F1 and by the absence of differentiation markers NR2F2, SOX17, AFP and ANPEP **(data accessible through NYSCF)**. Additionally, IHC staining for Oct4 and Tra-160 was performed to evaluate pluripotency **(Fig 2, for male lines).** Differentiation potency was assessed by in vitro embryoid body (EB)-based differentiation followed by gene expression analysis using Nanostring^57,58^ of genes expressed in the germ-layers **(data accessible through NYSCF)**. The absence of mycoplasma was confirmed with a biochemical enzyme assay. We confirmed that the identity of the gene-corrected lines matched the parental line by SNPTrace genotyping analysis^59^. Interestingly, we have not identified any clone that contained on-target effects in the targeted exon when correcting R83C or on-target effects in the neighboring exon when targeting the splice-site mutant. We found 10% on-target effects in the intron which are unlikely to affect the NAA10 protein. For the female line, we had a similar editing efficiency to correct R83C to the male line (2% and 1% positive clones); interestingly, the generation of a homozygous R83C clones was more efficient (42%), which might suggest that there is some selective advantage in cell culture to having the R83C/R83C genotype. We have not identified InDels on both alleles in any clone, one allele was always either WT or pathogenic variant. We conclude that total loss of NAA10 in hIPSCs leads to apoptosis, thereby no clones with InDels leading to a frameshift (early termination) in the male clones or in both alleles of the female clones were recovered **(Fig 4).** Processes for characterization and validation of cells lines can be seen in **Supplementary Table 2** (data accessible through NYSCF). The specific primers used for sgRNA, PCR, and ssODN can be found in **Supplementary Table 3**.

**Figure 3.**
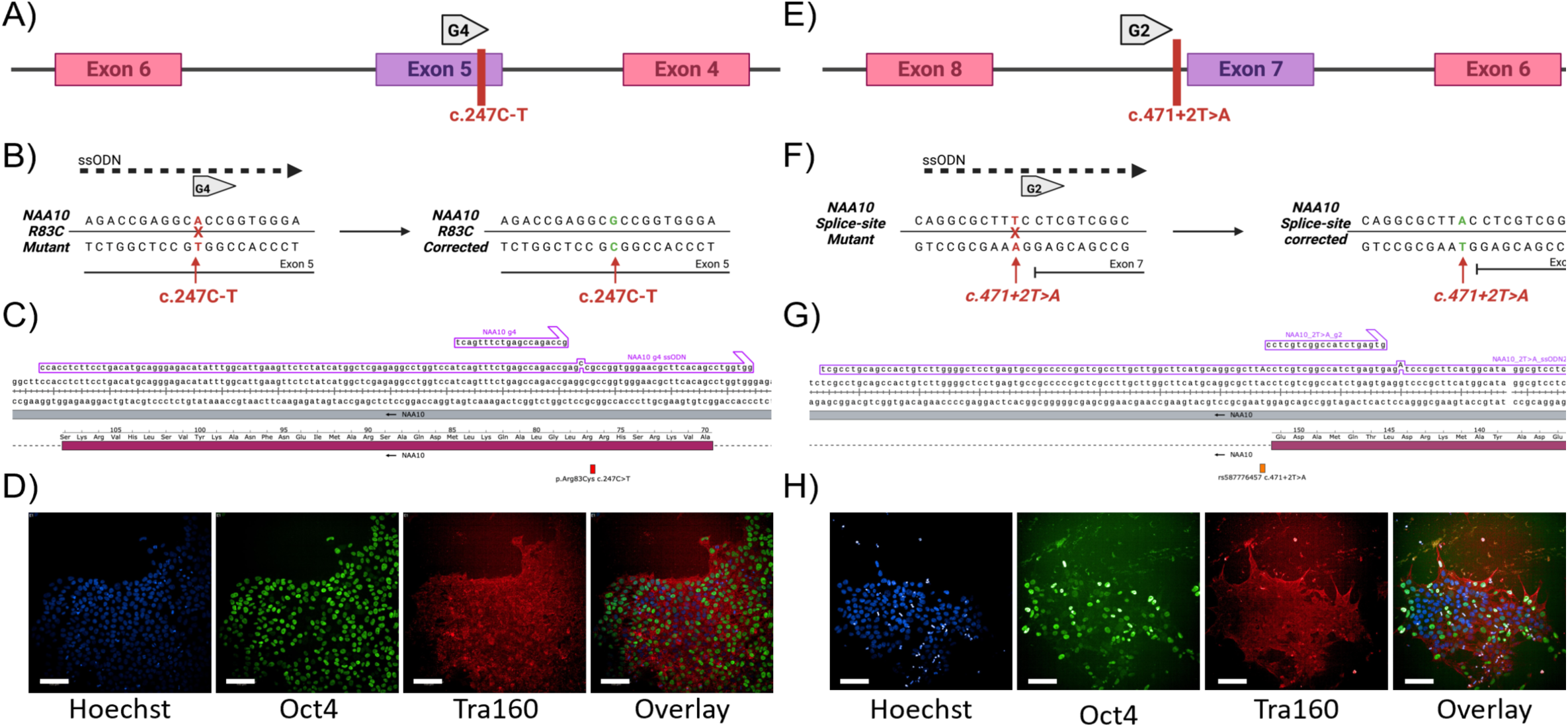
Generation of NAA10 corrected isogenic iPSC clones. A)+B) Strategy of editing NAA10 R83C. C) Detailed sequence and alignment of guide and ssODN. D) IHC for Oct4, Tra-160 and Hoechst as nuclear marker. E)+F) Strategy of editing NAA10 c.471+2T splice site correction. G) Detailed sequence and alignment of guide and ssODN. H) IHC for Oct4, Tra-160 and Hoechst as nuclear marker.

**Figure 4.**
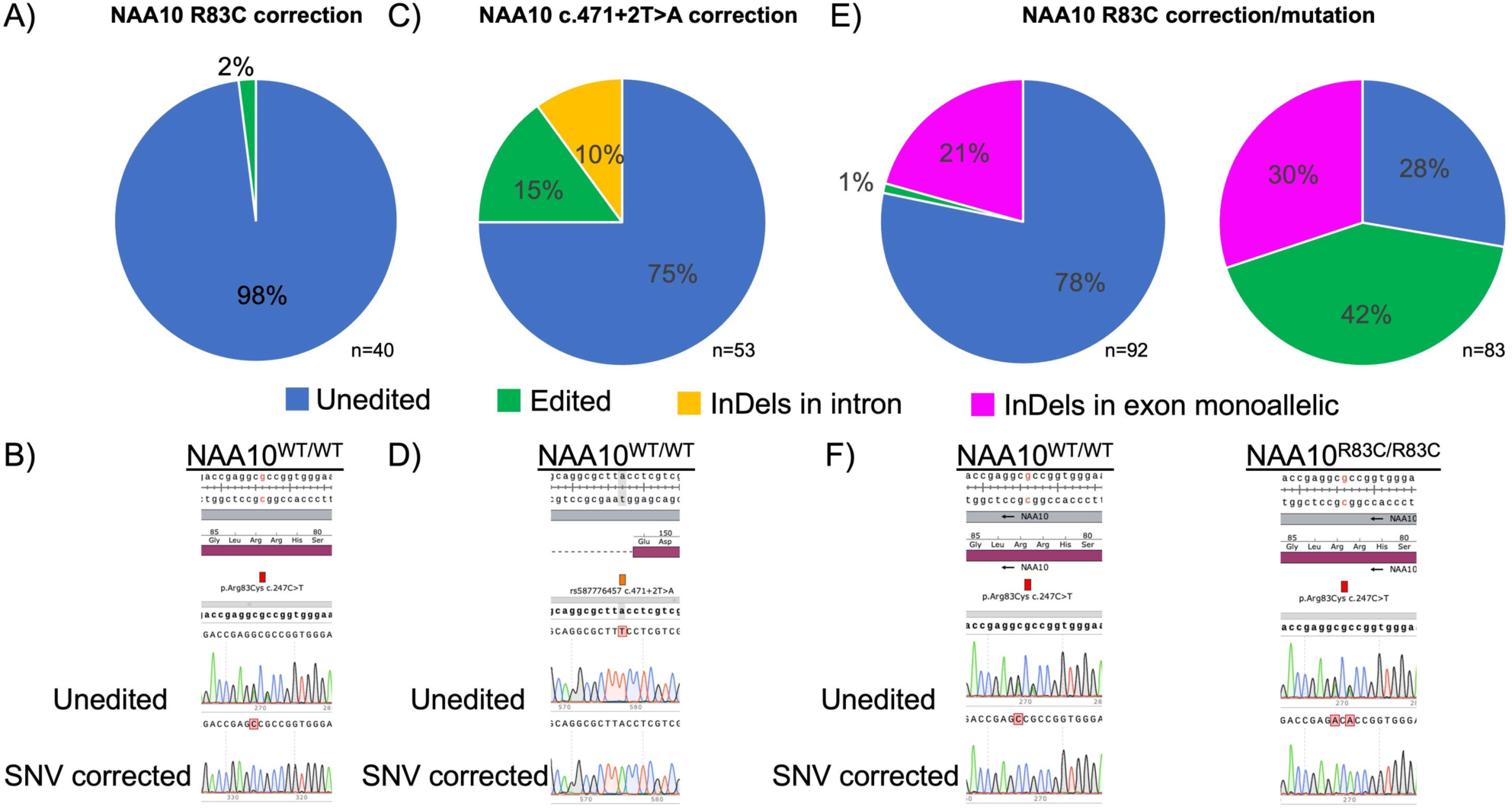
Outcome of gene editing. A) Analysis of all generated clones for the male NAA10 R83C line BR0010. B) Sequencing result of the parental unedited line and the positive clones. C) Analysis of all generated clones for NAA10c. 471+2T splice site pathogenic variant in the male line BR0017. D) Sequencing result of the parental unedited line and the positive clones. E) Analysis of all generated clones to generate a homozygous WT clone from the female line BR0015 (left) and a homozygous R83C clones (right) F) Left: Sequencing result for the parental unedited line and corrected clone for BR0015. Right: Sequencing result for the clone NAA10 R83C/R83C homozygous of the same BR0015 female line

**Table 2.**
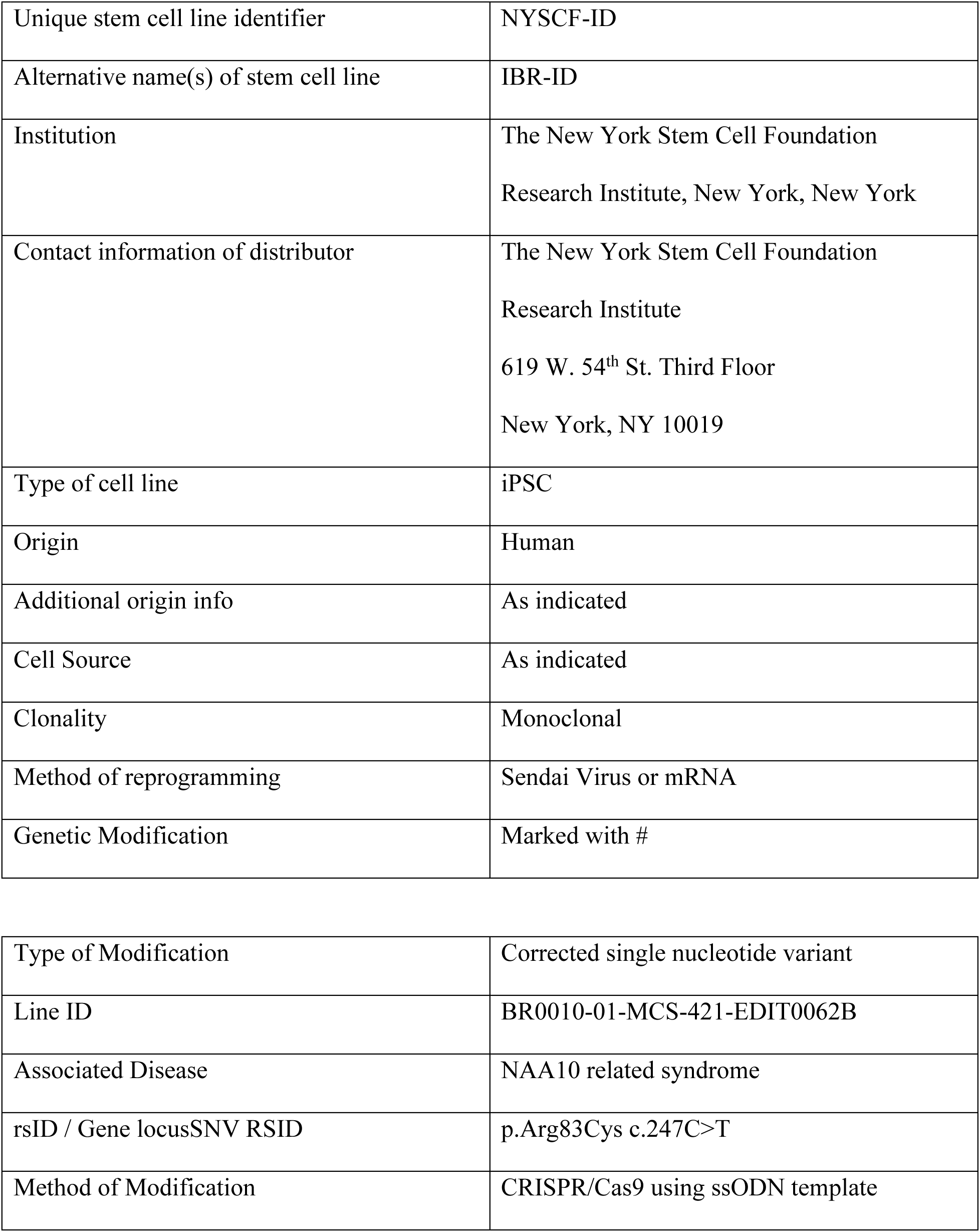
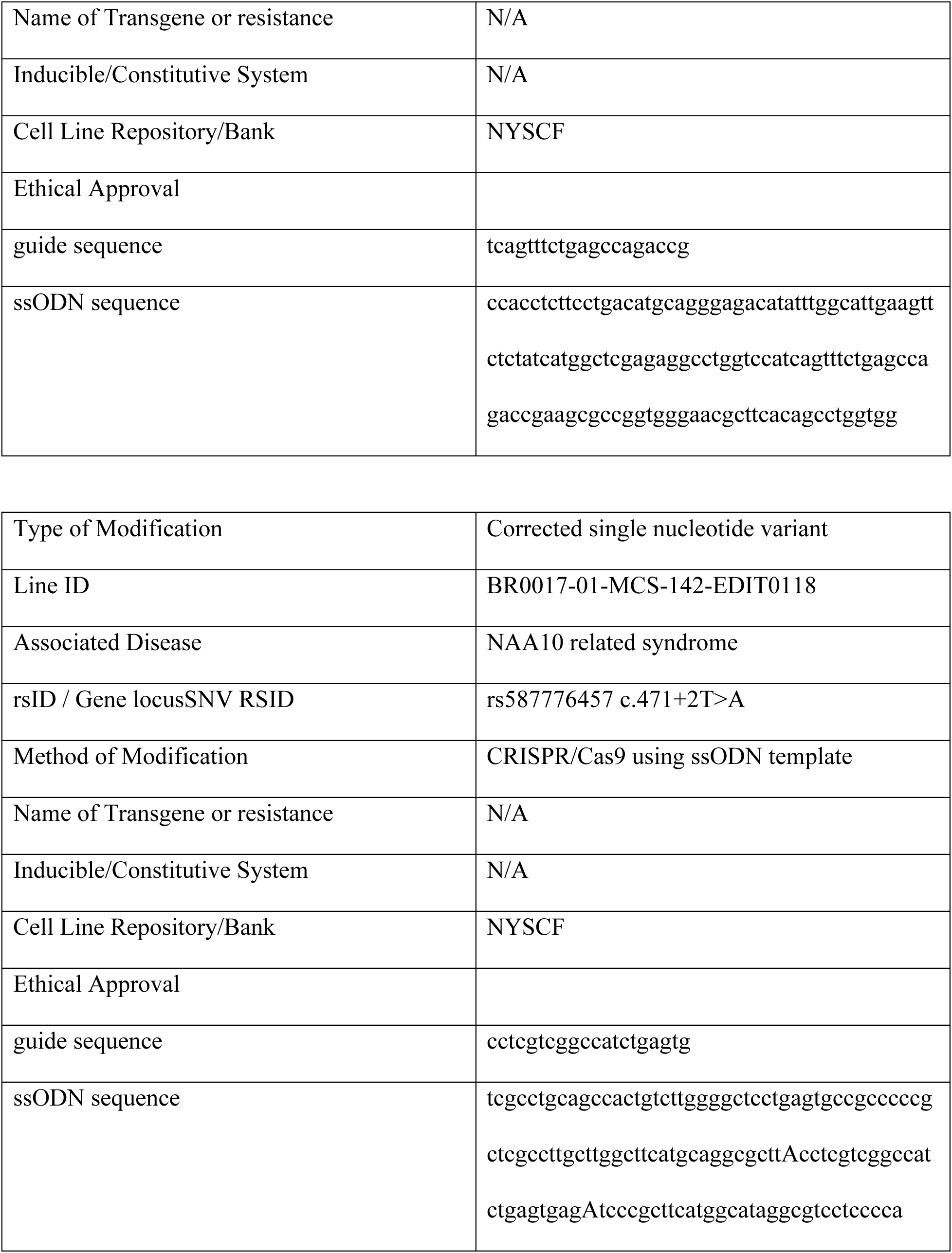
Characterization and validation of NAA10-corrected iPSC lines post ssODNs with Cas9-RNP correction.

### 2.4. Automated differentiation of iPSC derived neuronal progenitor cells (NPC’s)

Neuronal progenitor cells (NPC’s) were differentiated from iPSC cell lines using the high throughput automated differentiation (n=6) on the NYSCF Global Stem Cell Array® platform^58^ based on the dual SMAD inhibition protocol^60^ (**Fig 5**). Representative immunocytochemistry images of Neural progenitor cells (NPC) stained at Day 7 are shown in **Fig 6**.

**Figure 5.**
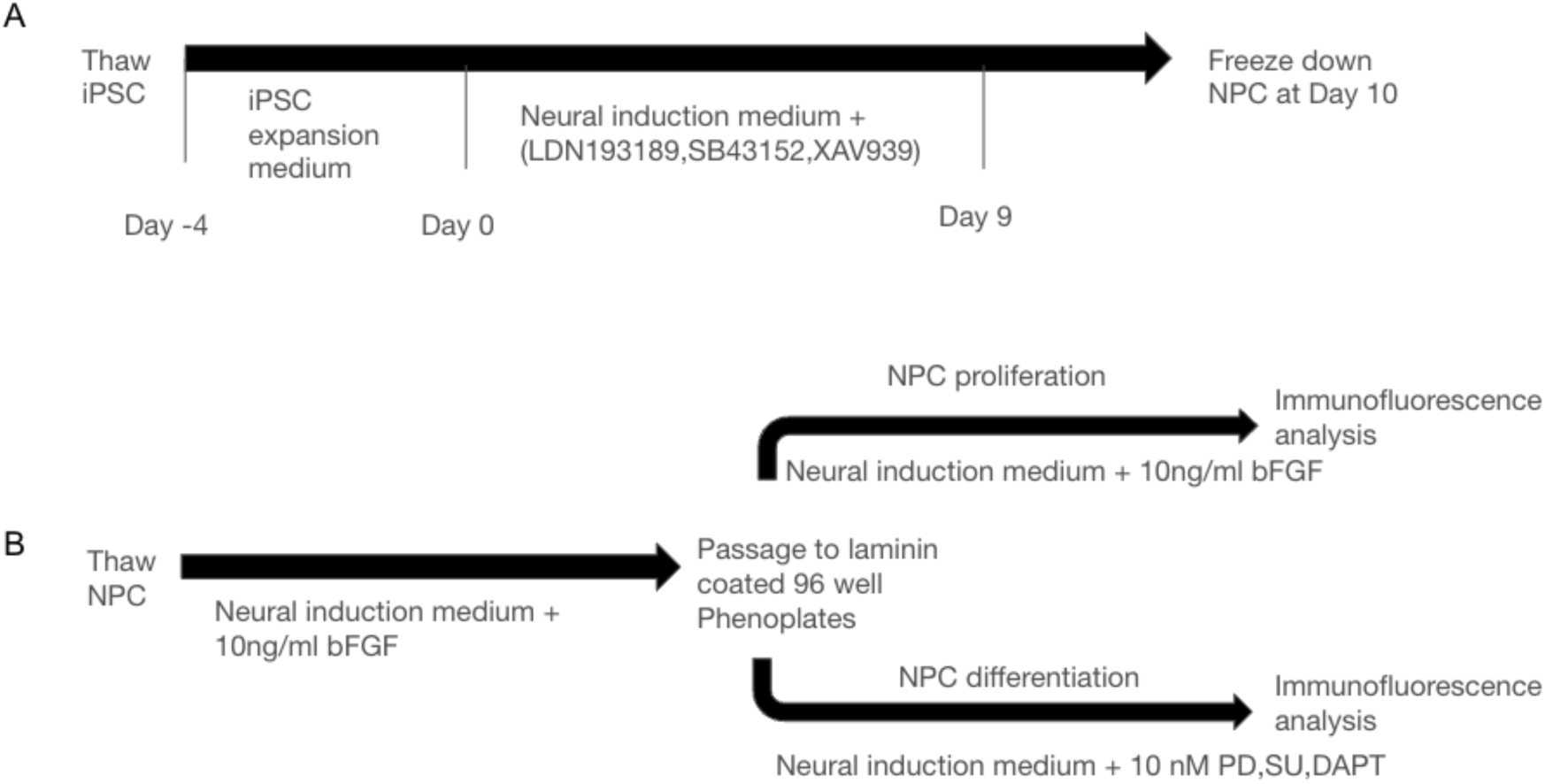
**A)** Schematic representation showing the differentiation timeline of neuronal progenitor cells (NPC’s) from the human induced pluripotent stem cells (hiPSC’s) with the media and the factors used for differentiation. **B)** Schematic representation showing the quality control (QC) assay of frozen NPC lines.

**Figure 6.**
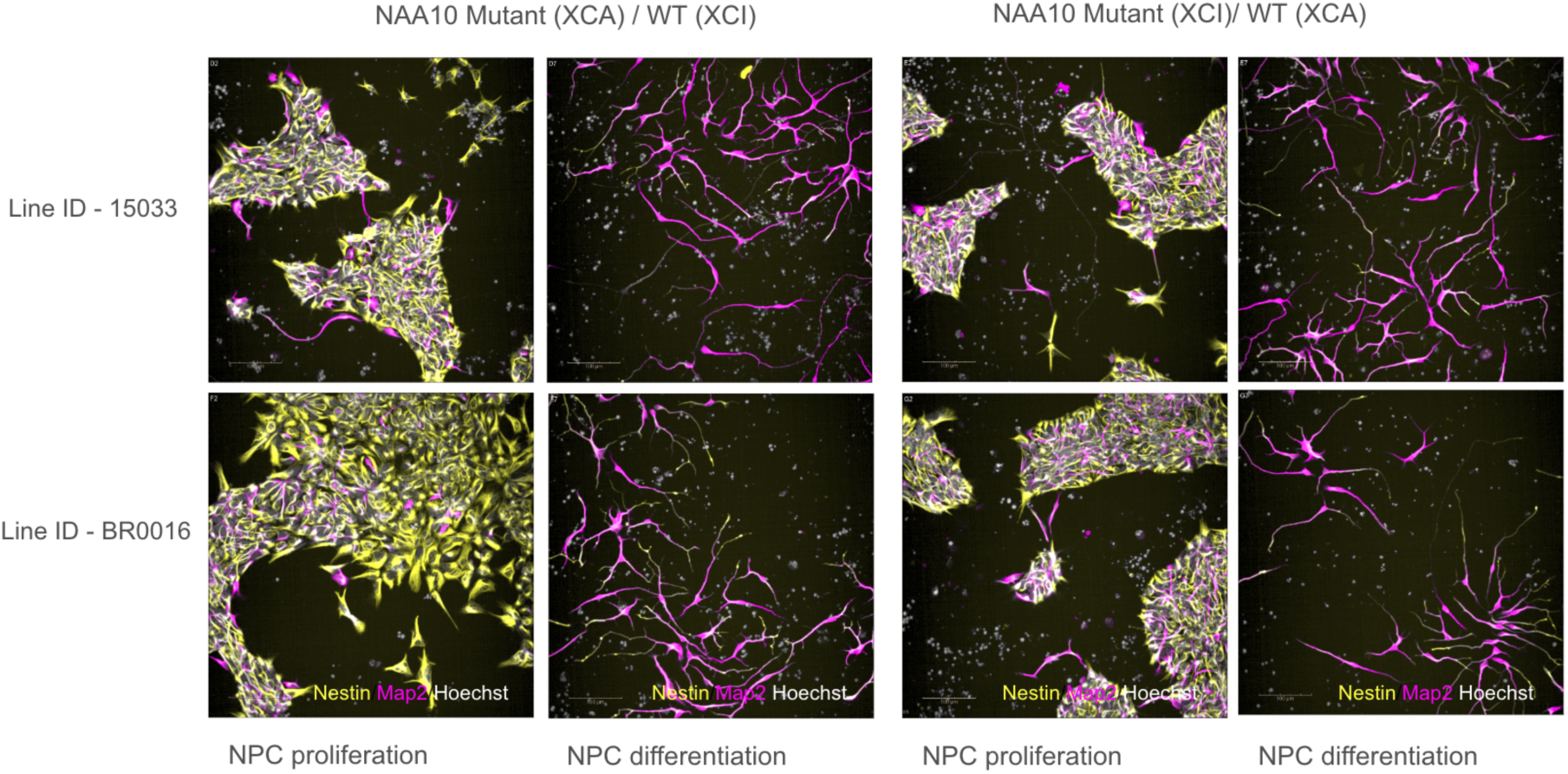
Immunocytochemistry images of Neural progenitor cells (NPC) stained at Day 7 and imaged at 20X magnification. Markers used for staining : Nestin(yellow), Map2 (pink), Hoechst (white). Scale bar - 100μM Row 1: - NPC’s for line ID 15033 supplemented with Neural induction medium + basic fibroblast growth factor (10ng/ml) showing NPC proliferation and supplemented with Neural induction medium+10 μM of PD0325901, SU5402 and DAPT showing NPC differentiation. Row 2: - NPC’s for line ID BR0016 supplemented with Neural induction medium + basic fibroblast growth factor (10ng/ml) showing NPC proliferation and supplemented with Neural induction medium+10 μM of PD0325901, SU5402 and DAPT showing NPC differentiation.

### 2.5. Validating the iPS lines and making cardiomyocytes

As further demonstration of the utility of this repository, a collaborator at the University of Utah (M.Y. and C.M.) received three isogenic pairs of male iPSC lines. Sanger sequencing confirmed the correct genotypes for these lines, followed by differentiation of the lines to cardiomyocytes, along with initial mass spectrometry-based proteomic analyses for one pair of iPSCs, which further confirmed differences in protein expression between the wild type and Arg83Cys mutant male BR0010 line (**Fig 7**).

**Figure 7.**
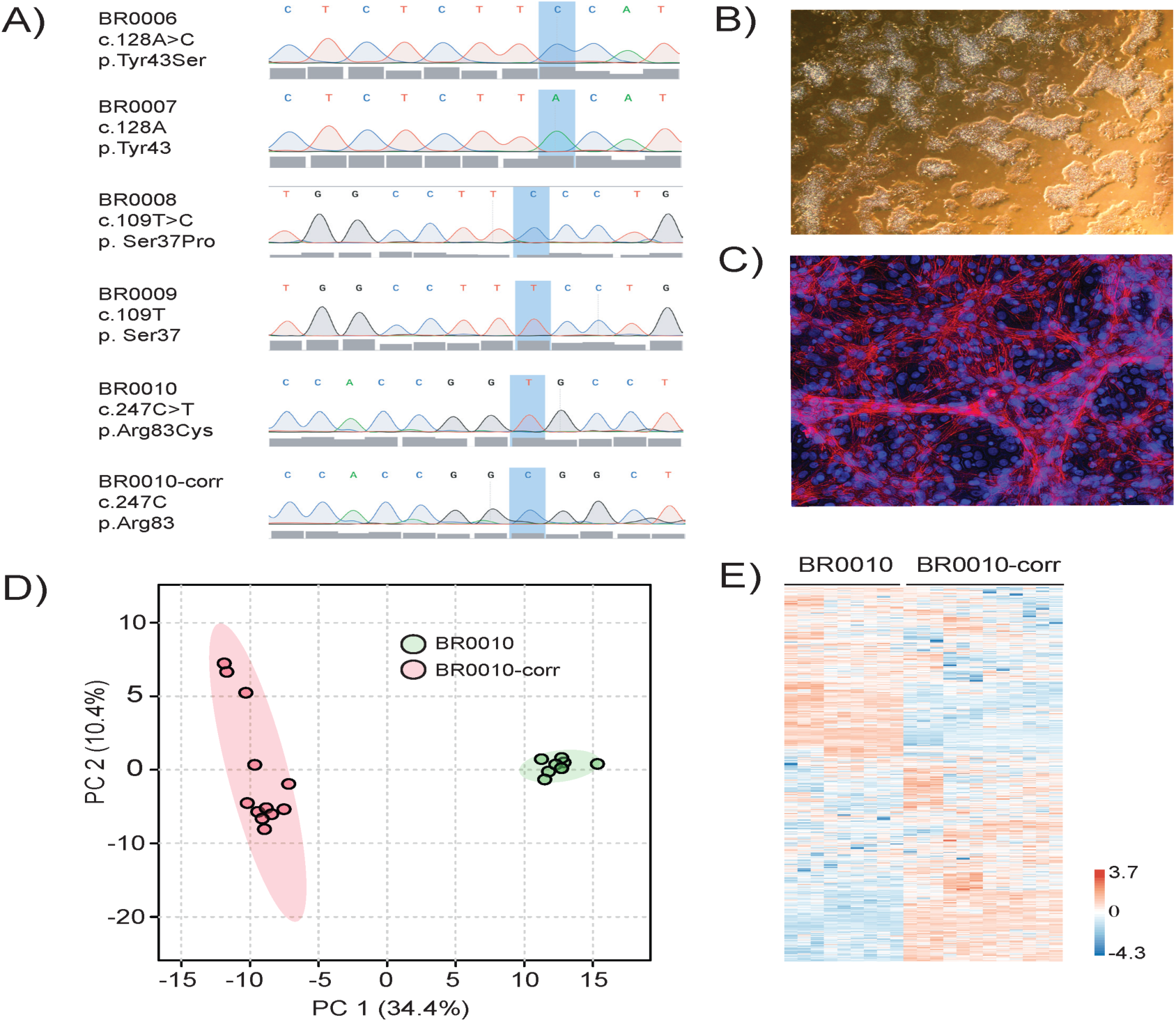
Validation of three male Ogden syndrome (OS) isogenic iPSC lines and Characterization of one male OS isogenic pair. Each line with the respective mutation or correction was verified by amplifying the region of interest and sequenced using primers. SnapGene was used to render the **A)** sequence chromatograms from the three isogenic Ogden syndrome lines: BR00006(Y43S), BR0007(Y43corrected), BR0008(S37P), BR0009(S37corrected), and BR0010(R83C), BR0010(R83corrected). The mutations and corrections of interest are highlighted in blue. Once iPSC lines were verified, stem cells were cultured and differentiated into cardiomyocytes (CMs). Representative images of BR0010[R83C] stem cells and CMs are shown in **panel B and C**, respectively. Day 14 post-differentiated CMs were stained with cardiac troponin T (red) and a nuclear Hoest dye (blue). In addition, day 14 post-differentiated CMs were lysed and prepared for LC-MS/MS. **D)** Principal component analysis of BR0010(R83C) and BR0010(R83corrected) cardiomyocytes, where each circle represents the combined abundance data for all proteins from a biological sample. The green circles within the PCA plot are the BR0010 line, and the red circles represent the corrected lines. **E)** Heatmap of all proteins identified in the BR0010 corrected and uncorrected line, based on log2 values based on normalized intensities.

## 3. Discussion

Previous studies with Ogden Syndrome derived iPSCs were used to characterize the long-QT phenotype present in two families with the p.S37P or p.Y43S disease causing variants^33^. With the establishment of a standardized protocol for the generation of iPSCs, future results can be cross validated in a manner that would minimize extraneous cell manufacturing processes and also provide for comparisons between iPSC lines with different disease-causing variants to better understand the electrophysiological aberrations causing the diseased phenotype. Furthermore, we used currently available protocols for inducing differentiation of iPSCs into neural progenitor cells^61^ and contractile myocytes^62^, thus further demonstrating the utility of these lines. Future investigation can involve utilizing these methods with the *NAA10* or *NAA15* iPSC lines to develop organoids for the creation of motor and higher order circuit models to better understand the physiological changes caused by each *NAA10* variant. Additionally, the development of novel therapeutics can be researched using these stem cell lines with methods of inducing X-chromosome reactivation^63,64^. Similar studies have been performed successfully with the recovery of FMR1 gene function in iPSC lines that model Fragile X syndrome^65^.

Our inability to recover any clones with InDels leading to frameshift in NAA10 **(Fig 4)** is consistent with the fact that *NAA10* was identified in screens for essential genes in human cell lines^66,67^. Unlike the situation in mice^68^, there is no currently known paralogue of *NAA10*, other than *NAA11* expressed in testicular and placental tissues^69^. That paper specifically looked at the lack of expression of *NAA11* in HeLa and HEK 293 human cells, where they used methylation-specific polymerase chain reaction and bisulfite sequencing to show that the absence of *NAA11* expression correlated with hypermethylation of the CpG island located at the proximal promoter of *NAA11*. They showed that a cloned *NAA11* gene promoter fragment was active when introduced into non *NAA11*-expressing human cells and its promoter activity was lost upon in vitro DNA methylation^69^. *NAA11* expression is therefore tissue-specific and is epigenetically regulated by DNA methylation. It is possible that *NAA11* expression could be re-activated during the course of attempting to knock out *NAA10* in human cells, such as iPS cells, but our inability to recover any indels in *NAA10* seems to indicate that this did not readily occur.

The original Ogden syndrome family (Ser37Pro) was reported as having four carrier women in it, who did not have any obvious cognitive phenotypes^26^. Recent testing with Vineland-3^70^ showed that two of these carrier women (I-2 and II-2 in the pedigree^39^) scored in the average range, with Vineland-3 ABC standard scores of 112 and 95, respectively. We previously published using a customized assay^87^, that the DNA isolated from blood from the carrier women in that family showed extreme X-chromosome skewing toward the wild-type (WT) allele, at close to 90% or higher. This skewing might also be toward the WT allele in the brains of these women, perhaps helping to explain why they are cognitively normal. The situation is quite different with other females with different pathogenic variants in *NAA10*, including Arg83Cys, where we have shown that such women are severely cognitively impaired^37^. It is possible that this missense variant is somehow much more deleterious toward NatA function, although *in vitro* assays using recombinantly expressed NAA10, NAA15 and HYPK gave inconsistent results, with NAA10 being more enzymatically active with Arg83Cys and severely impaired with Ser37Pro in the absence of HYPK, but at about the same level of reduced activity for both pathogenic variants in the presence of HYPK^28^. We have already written about the extensive limitations of such *in vitro* studies^29^, and the next step could include proteome-wide analyses of amino-terminal acetylation and protein expression levels in various cell types differentiated from the iPSCs, as we already performed in patient-derived skin fibroblasts, lymphoblastoid cells lines, and HeLa cells with knockdown of NatA^39^. This was also recently done for iPSCs with heterozygous loss of function, compound heterozygous, and missense residues (R276W) in *NAA15* introduced into the iPSC line (personal genome project 1) using CRISPR/Cas9 ^54^.

In relation to the different cognitive presentation for the carrier women with Ser37Pro and the other affected females, we endeavored to produce iPSC lines from multiple individuals with the same exact pathogenic variants, including five with Arg83Cys and three with Phe128Leu (**Table 1**). We have not yet made any iPSCs from the carrier women with Ser37Pro, but this might be something for future work, as it remains remarkable that they have no major phenotype. It is interesting that 4/5 of the Arg83Cys iPSC lines skewed toward WT on Xa in the primary sample (ranging from 84.5% to 89.9%), except for BR0016 (50.5%), which was the only primary sample that came from skin fibroblasts, as all other primary samples were from peripheral blood mononuclear cells (PBMCs). This led to difficulty with isolating clones with Arg83Cys on Xa, where we could only achieve this for BR0011 and BR0016. However, for Phe128Leu, 2/3 of the cell lines skewed toward pathogenic variant on Xa, for unknown reasons, leading to the isolation of only clones with pathogenic variant on Xa. As things currently stand, it is not known why the carrier females with Ser37Pro pathogenic variants are cognitively normal, whereas females with different pathogenic variants are severely affected. The iPSCs that we have created that are “isogenic” with mutant on Xa and/or wild type on Xa will enable further studies on the mechanism of how X-chromosome Inactivation (XCI) can have major effects on the outcome of disease, and this could have broader implications for other X-linked diseases^71–74^.

Ultimately, the major purpose of this article is to demonstrate our current pipeline for generating iPSC stem cell lines from human donors with NAA10- or NAA15-related neurodevelopmental syndromes and to provide a point of contact for collaborators should they be interested in ordering their own set of *NAA10* or *NAA15* iPSC cell lines for use in experimentation. Interested parties should reach out to the corresponding author G.J.L. or New York Stem Cell Foundation for requests. This repository is meant to facilitate new work by other groups on Ogden Syndrome.

## 4. Materials and Methods

### 4.1. Reprogramming [New York Stem Cell Foundation (NYSCF)]

For iPSC generation from peripheral blood mononuclear cells (PBMCs), cells are reprogrammed using Sendai virus mediated delivery of reprogramming factors using the NYSCF Global Stem Cell Array® (TGSCA™), a fully automated, robotic system that ensures high quality and decreases technical sources of variability^58^. Using automated protocols to count and passage cells, cells are transferred into 96 well plates at specified densities. Using automated transfection methods, Sendai virus containing the reprogramming factors hKLF4, Oct4,Sox2:hMyc:hKlf4 (Cytotune 2.0, Thermo Fisher) are added to the cells at a multiplicity of infection (MOI) of 5:5:3. Cells are cultured initially in Blood Reprogramming Media (Complete Stempro34 supplemented with Glutamax and SCF (200 ng/μL), FLT3 (200 ng/μL), IL-3 (40 ng/μL), and IL-6 (40 ng/μL) (all Thermo Fisher)) before being transferred to Freedom media (A14577SA, Thermo Fisher). 10-20 days post reprogramming, colonies are identified using live Tra-1-60 Cell surface marker staining (R&D Systems). Cells are harvested into intermediate stocks before entering an enrichment/monoclonalization step. We utilize automated methods to prepare samples for fluorescence-activated cell sorting (FACS) enrichment, allowing the depletion of non-iPSC cells (Tra-1-60-) and the seeding of single cells for monoclonal outgrowth. During this time, cells are fed with Freedom media. All dissociation steps take place using Accutase (Thermo Fisher), with the cell culture media supplemented with either 1 μM Thiazovivin (Sigma) or CloneR (Stemcell Technologies). Monoclonal lines are derived following ideal morphological selection of clonal outgrowths using our established machine learning framework called Monoqlo℠, which automatically detects clonalized cell colonies and assesses clonality from daily imaging data before further intermediate stocks of the monoclonal lines are frozen. Two iPSC lines are generated per donor, with one brought forward for distribution and the other stored as a backup. Following the completion of iPSC monoclonalization, cells are entered into an expansion workflow that sees the creation of a distributable inventory of iPSCs and material for quality control (QC).

For iPSC generation from skin fibroblasts, cells are reprogrammed using non-modified RNA and microRNA technology on the TGSCA. Cells are seeded onto a pre-warmed i-Matrix coated 24-well plate at a density of 30,000 cells/well. After preparing the mRNA reprogramming cocktail (Stemgent 00-0076), plates are fed with Nutristem (Stemgent) and transferred to a hypoxic incubator for pre-conditioning. Later that day, at least 6 hours after the Nutristem feed, the transfection master mix is prepared and added to each well for transfection. Cells are fed with Nutristem each morning and transfected each afternoon for 10 days. Cells then undergo a negative sort using the Miltenyi Biotec MultiMACS cell separation system anti-fibroblast magnetic beads. iPSC colonies are identified and a live cell Tra-1-60 antibody is used to confirm pluripotency through daily imaging. Positive colonies are consolidated into intermediate stocks and then entered into an expansion workflow to create a distributable inventory of iPSCs and material for QC.

All reprogrammed iPSC lines are subjected to quality control assays using automated workflows already established in the NYSCF GSCA, including post-thaw cell recovery, sterility, mycoplasma test, karyotype, identity test, pluripotency profile, Sendai transgene exclusion, and differentiation capacity **(Supplementary Table 2)**. All cells are frozen in 2D barcoded Matrix Tubes (Thermo Fisher) **(Supplementary File 1)**. Using such tubes allows barcode tracking of all samples using our proprietary NYSCF Laboratory Information Management System (LIMS) system. All samples, cell culture plates, and reagents are barcoded and tracked through this system.

### 4.2. X-chromosome Screening [New York Stem Cell Foundation (NYSCF)]

RNA was extracted from frozen primary samples, polyclonal iPSCs or monoclonal iPSCs with RNeasy Mini Kit (Cat# 74104, QIAGEN). RNA concentration was measured with Nanodrop (Thermo Fisher). Reverse transcription was performed with SuperScript™ III First-Strand Synthesis SuperMix (Invitrogen) and PCR was performed with the AmpliTaq Gold™ 360 Master Mix (Applied Biosystems) using primers shown in **Table 3**. PCR amplicons of primary and iPSC pool samples were submitted for Next Generation Sequencing to assess X-chromosome activation status. PCR amplicons from monoclonal iPSCs were sent for Sanger sequencing (Genewiz). Sequencing results were analyzed using Snapgene.

**Table 3.**
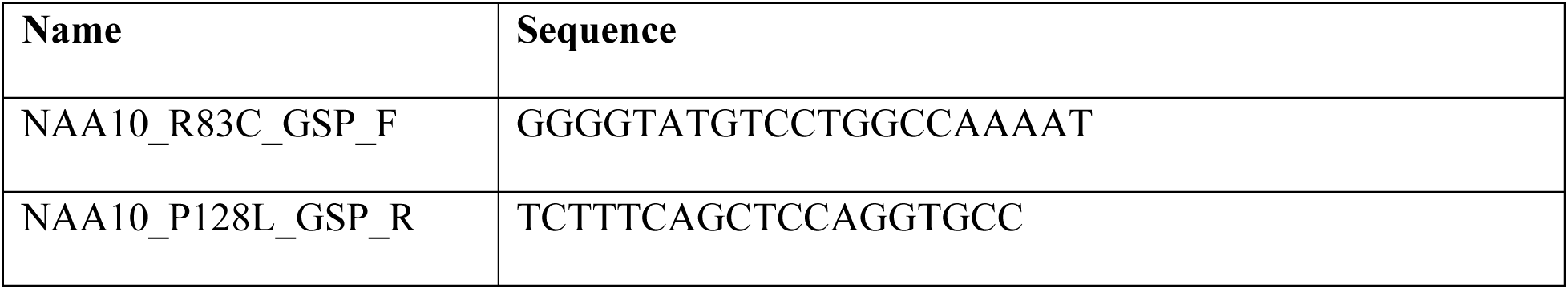
Primer sequences for PCR.

### 4.3. iPSC culture [New York Stem Cell Foundation (NYSCF)]

Human iPSCs maintained on Cultrex (CTX, R&D Systems) in PSC Freedom Media (FRD1, Thermo Fisher Custom) were passaged every 4-5 days using Accutase (Thermo Fisher) in the presence of 1uM Thiazovivin (THZ, Sigma-Aldrich).

### 4.4. sgRNA and ssODN Design [New York Stem Cell Foundation (NYSCF)]

CRISPR^75^ was used to design guides. Single stranded ODNs were designed by going 91 bps downstream from the cutting side of the guide and 36 bps upstream. Blocking mutations were added to the ssODNs so that either the protospacer adjacent motif (PAM) site was destroyed or 2 mismatches in the middle of the guides would appear. Those blocking mutations do not lead to a change in the amino acid chain.

### 4.5. CRISPR/Cas9-mediated gene editing Transfection [New York Stem Cell Foundation (NYSCF)]

Human iPCs were dissociated using Accutase and 1.6 x10^5^ cells were seeded onto a 96-well round bottom plate in FRD1 containing CloneR from a 12-well source plate containing these hiPSCs in log phase. Transfection cocktails were prepared using Lonza-P3 Primary Cell NucleofectorTM X Kit. The transfection cocktail contained P3-buffer+supplement as specified by Lonza, 2ug Alt-R™ S.p. Cas9 Nuclease V3, 1.9 ug Alt-R CRISPR-Cas9 sgRNA (IDT) and 40uM Alt-R HDR Donor Oligo (IDT). The passaged cells were pelleted and transfection cocktail was used to resuspend them to create a 20ul suspension. Electroporation was carried out in 20 µL 16-well Nucleocuvette^TM^ Strip format using the CA-137 program. Post nucleofection cells were plated in triplicates on a CTX pre-coated 97-well corning flat bottom plate. They were subjected to cold shock at 32C for 24hrs followed by a media exchange with FRD1 media. 72hrs after transfection efficiency was checked using PCR, sanger sequence and Sythego’s ICE analysis to determine for monoclonalization.

### 4.6. Monoclonalization [New York Stem Cell Foundation (NYSCF)]

Transfected iPSCs with successful ICE-analysis score were single cell sorted into 96 well plates using a Benchtop Microfluidic Cell Sorter (BDFACSAria III Cell Sorter). Plates were fed daily with FRD1 and scanned every night on a Celigo Image Cytometer (Nexcelom Bioscience). After 10 days monoclonal colonies were consolidated and transferred into a new 96 well plate. Wells were transferred when reaching 80-100% confluency for freeze backs and sequencing analysis.

### 4.7. Sanger sequencing of monoclonal wells [New York Stem Cell Foundation (NYSCF)]

Quick extract gDNA template was prepared by depositing 5.0 x 10^4^ cells into a 96 well hard-shell PCR plate (Bio-Rad). The plate was centrifuged, media aspirated and 30uL of QuickExtract™ DNA Extraction Solution (Lucigen) added to the wells. The sealed PCR plate was then run through the QuickExtract heating cycle as per the manufacturer’s instructions.

### 4.8. Mycoplasma & Sterility [New York Stem Cell Foundation (NYSCF)]

In order to ensure that the samples arrived without mycoplasma contamination, and none was inadvertently introduced during production, media was collected for mycoplasma testing at two points across the process: after the first MSC passage and during the first passage of iPSC expansion post thaw. Testing for mycoplasma contamination was done robotically with the MycoAlert Mycoplasma Detection kit mycoplasma luminescent assay (Lonza, #LT107-318) and the accompanying MycoAlert Assay Control Set (Lonza, #LT07-518) and read on an integrated Synergy HT plate reader (BioTek). Non-mycoplasma contamination was assessed via incubation of supernatant media, from the first passage of iPSCs post thaw, with Tryptic Soy Broth (Hardy Diagnostics). Absorbance reads were conducted at 0, +24, +72, and +168 hours after sterility plate creation, to assess any growth. Additionally, iPSC cultures were monitored using Nexcelom Celigo Image scans daily.

### 4.9. Karyotyping [New York Stem Cell Foundation (NYSCF)]

Karyotype analysis was performed at passage 13 using the Illumina Core-Exome24 or Global Screening Array genotyping chip, with data analyzed via GenomStudio (Illumina) and the CNV analysis, cnvPartition 3.2.0. The absence of major (>2.5 Mb) insertions, deletions, or chromosomal aberrations was used to confirm a normal karyotype.

### 4.10. Identity [New York Stem Cell Foundation (NYSCF)]

DNA was extracted from both the primary sample and the iPSCs prior to cryopreservation at the end of expansion. It was extracted using an epMotion liquid handler (Eppendorf) and ReliaPrep 96 gDNA Miniprep HT System (Promega). The DNA was tested on the Fluidigm Juno system using the SNPTrace platform to analyze 96 unique SNPs. The line passed the assay if the DNA from the primary sample and iPSCs matched with high confidence, minimum of 92 out of 96 SNP match.

### 4.11. Pluripotency Expression Profile [New York Stem Cell Foundation (NYSCF)]

RNA was extracted from the iPSCs prior to cryopreservation at the end of expansion. The RNA was assayed on the Nanostring nCounter Flex system using Nanostring’s Patented Molecular Barcoding System to tag and count un-amplified RNA Targets^58,76^ The data was normalized against a pre-established panel of human embryonic stem cell (hESC) lines. The line passed the assay if the iPSCs showed expression of pluripotency-associated genes and absence of spontaneous differentiation-associated markers.

### 4.12. Shutoff of Sendai Transgene [New York Stem Cell Foundation (NYSCF)]

RNA was extracted from the iPSCs at the end of expansion. The RNA was assayed on the Nanostring nCounter Flex system using Nanostring’s Patented Molecular Barcoding System to tag and count un-amplified Sendai virus backbone RNA targets. The line passed the assay if there was low to no expression of the Sendai virus backbone.

### 4.13. Differentiation Capacity [New York Stem Cell Foundation (NYSCF)]

iPSCs were passaged to an Elplasia 96 well microcavity plate (Corning) for embryoid bodies (EBs) to spontaneously form over 16 days. EBs were collected, lysed, and assayed on the Nanostring using Nanostring’s Patented Molecular Barcoding System to tag and count un-amplified RNA Targets. The data was compared against a pre-established panel of human embryonic stem cell (hESC) lines spontaneously formed into EBs and analyzed using custom scripts based on previously published data scorecard analysis^57^. The line passed the assay if the iPSCs displayed levels of gene expression for germ layer markers consistent with the hESC-derived EBs. The score for each of the three germ layers was provided in the certificate of analysis **(Supplemental File 2)**.

### 4.14. Post-Thaw Viable Cell Recovery [New York Stem Cell Foundation (NYSCF)]

After freezing down iPSCs into final Repository tubes, one tube was thawed directly into one well of a 12-well plate using recommended culture conditions, StemFlex Media (Gibco) and Cultrex (R&D systems) followed by daily feeds; CloneR is used in the initial 24h of thaw. This replicated the thawing protocol described in the NYSCF SOP **(Supplemental File 3)**. The line passed the assay if the cells reached greater than 50% confluency within 10 days post thaw, without any indication of spontaneous differentiation or contamination.

### 4.15. Immunohistochemistry staining [New York Stem Cell Foundation (NYSCF)]

iPSCs were passaged onto a 96 well CCU plate and fed for 3 days, then fixed in 4% PFA for 10 min, permeabilized in PBS, 0.2% Triton-X 100, 1% H-FBS for 30 min. Fixed-perm cells were washed with PBS+FBS and blocked for 15 min with PBS and 5% FBS. A 2x antibody mix containing Anti-TRA-1-60-PE (Miltenyi Biotec, Cat.120-007-552) Alexafluor 488 Oct4 (BD Bioscience, Cat.560253) was added to the blocking solution and incubated at 4C over night. iPSCs were washed with PBS three times. Hoechst was added at the second wash and incubated for 10 min. Cells were imagined on a Phenix. Specific components, reagents, and concentrations can be found in **Supplementary Table 4**.

### 4.16. Automated differentiation of iPSC derived neuronal progenitor cells (NPC’s) [New York Stem Cell Foundation (NYSCF)]

The iPSC lines were thawed in an expansion medium (Life Technologies custom media #A14577SA) supplemented with 10% CloneR (Stemcell Technologies, #05888) and coated on a Cultrex (R&D systems, #3434-010-02) coated Corning**™** (Fisher Scientific, #07-200-82) Costar**™**12 well plate. After allowing them to expand to about 80-90% confluence the cells were passaged into 4 Corning **12** well plates seeding them at a density of 150,000 cells/cm^2^. 24 hours after the passage the cells were changed to the Neuronal induction medium (NIM) which consists of a 1:1 ratio of basal media containing DMEM/F12+Glutamax (Thermo Fisher, #10565042) and Neurobasal (Thermo Fisher, #21103049) supplemented with Glutamax (Thermo Fisher, #35050-061), B-27 with vitamin A (Thermo Fisher, #17504044) and N2 (Thermo Fisher, #17502-048). To induce differentiation the NIM media was supplemented with the small molecules LDN193189 (Sigma, #SML0559), SB431542 (Sigma, #S4317), XAV939 (Sigma, #X3004). The media exchange was performed daily for a duration of 10 days for the entire period of differentiation. On day 10 the cells were frozen down into matrix tubes at a density of 1 million cells per vial using the CryoStor® freezing medium (Stemcell Technologies, #100-1061) **(Figure 5A)**.

To perform the Quality control (QC) we used 2 PhenoPlate™ 96-well microplates (Revvity, #6055302) coated with 0.1% polyethylenimine (PEI) (Sigma, #408727) in 0.1M Borate buffer pH 8.4. After washing the PEI solution with water the PhenoPlates were coated with 10 µg/ml laminin solution (Thermo Fisher, #23017015). For the QC the cells were plated on Cultrex coated Corning**™**Costar**™**96 well plates (R&D systems, #07-200-90) in NIM+10 % CloneR. Media exchange was performed with NIM+ 10 ng/ml basic FGF (R&D systems, #233-FB-010) on Day3 after thaw allowing them to proliferate. Once the 96 wells were confluent with NPC they were passaged onto the 2 laminin coated PhenoPlates at variable seeding densities. One of the plates would be used to check for the NPC proliferation and the 2^nd^ plate would be used to check the neuronal induction of the NPC **(Figure 5B)**. For the NPC proliferation the media exchange was done using NIM+ 10 ng/ml basic FGF. For the neuronal induction the media exchange was performed using NIM+10uM PD0325901 (Sigma, #PZ0162),10uM SU5402 (Sigma, # SML0443) and 10uM DAPT (Sigma, # D5942). Media exchange was performed every other day for 7 days and to detect the presence of the neuronal markers Nestin and Map2 immunofluorescence assay was performed.

For immunofluorescence analysis by adding 32% paraformaldehyde (Electron Microscopy Sciences) directly to medium to a final concentration of 4% and incubated at room temp for 15 min. Cells were washed three times with HBSS (Thermo Fisher Scientific), stained overnight with mouse anti-Nestin 1:3,000 (Millipore, Cat.09-0024), chicken anti-MAP2 1:3,000 (Abcam, Cat.09-0006) in 5% normal goat serum (Jackson ImmunoResearch) in 0.1% Triton X-100 (Thermo Fisher Scientific) in HBSS. Primary antibodies were counterstained with goat anti-mouse Alexa Fluor 555 and goat anti-chicken Alexa Fluor 647 and 10 µg/ml Hoechst for 1 hour at room temp. Cells were washed three times with HBSS.

### 4.17. iPSC culture [University of Utah]

Induced pluripotent stem cell lines (iPSCs) from Ogden syndrome patients were received from New York Stem Cell Foundation. We also received 4 earlier passage lines directly from our collaborators at Stanford who had worked previously on this^33^. To further validate the iPSC lines and study Ogden syndrome, three male isogenic lines (6 lines total) were cultured in 6-well plates coated with vitronectin using E8 and StemFlex media. The 6 lines include: BR00006(Y43S), BR0007(Y43corrected), BR0008(S37P), BR0009(S37corrected), BR0010(R83C), and BR0010(R83corrected). Cells were grown to 60-80% confluency before differentiation or dissociation with EDTA to passage. Cell pellets for DNA extraction and karyotyping were also generated by dissociating cells with EDTA and centrifuging at 300 x g for 2 min prior to flash freezing and storing at −80°C.

### 4.18. Sanger sequencing validation of iPSC lines [University of Utah]

To verify the mutation or correction of interest **(Figure 7A)**, DNA was extracted from individual cell pellets using a DNA Mini Kit (Qiagen, Cat.51306). Primers were used to amplify the region of interest using a SimpliAmp Thermal Cycler (Thermo Fisher Scientific) and the PCR product was cleaned up using Genomic DNA Clean & Concentrator (Zymo Research, Cat.D4011). The purified PCR product was then sequenced at the University of Utah Genomics Core. The Naa10 forward primer: 5’-TCACCGCCGCCTTAGACTGA-3’ and reverse primer: 5’-ATAGCACCCCTCAGCATCCCCT-3’ were used to sequence BR0006/BR0007 isogenic lines (Y43S, c.128A>C) and BR0008/BR0009 isogenic lines (S37P, c.109T>C). To sequence the BR0010 isogenic lines (R83C, c.247C>T), the Naa10-R83C forward primer: 5’-GCATGTCCACTCTACAAATGGC-3’ and reverse primer: 5’-ATACTGCCTTGACGGGGGTC-3’ were used.

### 4.19. Mycoplasma & Sterility [University of Utah]

To ensure that the samples arrived without mycoplasma contamination, and none was inadvertently introduced during production, iPS cells were grown and DNA extracted as described above in the methods section 4.17 and 4.18. Each sample was then tested for mycoplasma using the Microsart AMP Mycoplasma kit (Sartorius, Cat.SMB95-1005) following the manufacturer’s instructions. PCR reactions were run and analyzed using QuantStudio 12K Flex and software at the Genomics Core of the University of Utah.

### 4.20. Differentiation of iPSC derived cardiomyocytes (iPSC-CMs) [University of Utah]

After sequencing and sterility were confirmed, the three isogenic male Ogden syndrome iPSC lines were differentiated into iPSC-cardiomyocytes (CMs) using a modified protocol from Burridge et al.^77^ In brief, iPSCs were cultured using E8 media in 6-well plates coated with vitronectin. When cells reached 60-80% confluence in 3-5 days after passaging, media was switched to CDM3 (RPMI1640 medium + GlutaMAX (Gibco, Cat.61870036), 250 µg/mL hAlb (Sigma, Cat.A9731-5G), 250 µg/mL BSA (Gibco, Cat.11020-021), and 215 µg/mL ascorbic acid (Sigma: A8960-5G)) as a basal media for cardiomyocyte (CM) differentiation. Media was changed with fresh CDM3 every day for at least 6 days. After day 6, media is changed at least every other day. In addition, CHIR99021(Selleckchem, Cat.S1263), Wnt-C59 (Selleckchem, Cat.S7037), and RevitaCell (Gibco, Cat.A2644501) were added to the CDM3 media to aid in CM differentiation. For days 0-1, CHIR99021 (3 µM) was added to each well to activate the Wnt pathway. For days 2-3, Wnt-C59 (2 µM) was added to each well to inhibit the Wnt pathway. For days 0-5, RevitaCell (0.23X) was added to each well to prevent cell death. Onset of beating is typically observed between days 8-10 post-differentiation.

### 4.21. Immunohistochemistry staining of iPSC-CMs [University of Utah]

iPSC-CMs were stained on day 14 post-differentiation using RV-C2 Troponin T, cardiac type^78^ (DSHB, RV-C2) and a nuclear Hoechst dye (Thermo Scientific, Cat.62249) **(Figure 7C)**. All steps of the immunostaining process were performed at room temperature. To prepare the cells for staining, media was aspirated from each well (from a 6-well plate), washed with 1 X DPBS, and fixed with 4% PFA (Thermo Scientific, Cat. 28906) for 10 minutes. The cells were washed with 1 X DPBS + 0.5% BSA between each subsequent step. Cells were then permeabilized with PBT for 30 minutes, followed by the primary antibody for 30 minutes and secondary antibody AF594 (Thermo Fisher Scientific, Cat. A21145) for 30 minutes. Finally, the cells were stained with a nuclear dye (Hoechst) for 10 minutes and stored in 1 X DPBS (protected from light). Images of iPSC-CMs were acquired using an ECHO revolve and EVOS M7000 microscope.

### 4.22. iPSC-CM cell lysis for mass spectrometry [University of Utah]

On day 14 post-differentiation, iPSC-CMs were rinsed with 1 X DPBS and dissociated with a cell scraper in RIPA buffer (Thermo Fisher Scientific, Cat.89900) supplemented with 1 X protease inhibitors (Thermo Fisher Scientific, Cat.78442). Cells were then added to a pre-chilled 1.5 mL Eppendorf tube with 0.1 mm and 0.5 mm glass beads and incubated on ice for 30 minutes. Cell lysis was performed by vortexing at high speed (7-8) for 10-minute intervals at 4°C, repeated four times. In between each interval, the sample tubes were incubated ice for 3-5 minutes. Once lysed, the samples were spun down in a cooled centrifuge (4°C) at max speed for 10 minutes. The soluble supernatant was then transferred to a new low protein binding microcentrifuge tube (Thermo Fisher Scientific, Cat.90410) and flash frozen/stored at −80°C.

### 4.23. Protein Digestion [University of Utah]

Ten microgram of lysate were added to 200 μL urea buffer (8M Urea, 0.1M Tris/HCl pH 8.5) and loaded into 30 KD Vivacon 500 filter units and centrifuged at 13,000g for 15 minutes, and then the concentrated protein was washed three times with urea buffer. The concentrate was alkylated with 50 mM iodoacetamide in urea buffer and incubated in the dark at room temperature for 20 minutes, followed by centrifugation for 15 minutes. The concentrate was washed twice with urea buffer and two washes with 50 mM ammonium bicarbonate. 10 μg of protein was subjected to trypsin digestion, added at a 1:40 enzyme ratio, and incubated for 18 hours at 37°C. The peptides were then collected by centrifugation at 13000g for 15 minutes. The filters were washed with 50 mM ammonium bicarbonate, and the wash was also collected by centrifuge at 13000g for 15 minutes. The collected peptides were acidified to 1% formic acid and placed into mass spectrometry vials for analysis.

### 4.24. Mass Spectrometry (MS) Analysis [University of Utah]

An isogenic pair (BR0010(R83C) and BR0010(R83corrected)) was chosen for MS analysis. BR0010(R83C) cardiomyocytes (passage (P) 15, 16, and 20) were grown on 3 different 6-well plates. Six biological replicates (one well per biological replicate) were used to analyze the proteome of BR0010(R83C). Similarly, BR0010(R83corrected) cardiomyocytes (P13 and 14) were grown on 2 different plates. Seven biological replicates were used to analyze the proteome of BR0010(R83corrected).

Tryptic peptides were analyzed as previously published^79–82^ by nanoflow LC-MS/MS on a Thermo Orbitrap Velos Pro interfaced with a Thermo EASY-nLC 1000 equipped with a reverse-phase column (75µm inner diameter, 360 µm OD, 15cm, Reprosil-Pur 120 C-18 AQUA 3µm particle size; ESI Solutions) and a flow rate of 400 nl/min. For peptide separation, a multi-step gradient was utilized from 98% Buffer A (0.1% formic acid, 5% DMSO) and 2% Buffer B (0.1% formic acid, 5% DMSO in acetonitrile) to 10% Buffer A and 90% Buffer B over 90 minutes. The spectra were acquired using Nth order double-play, data-dependent acquisition mode for fragmentation in the parent spectra’s top 20 most abundant ions. MS1 scans were acquired in the Orbitrap mass analyzer at a resolution of 30000. MS1 ions were fragmented by either collision-induced dissociation. Dynamic Exclusion was enabled to avoid multiple fragmentations of parent ions.

### 4.25. Mass Spectrometry Data Analysis [University of Utah]

The resulting spectra were analyzed using MaxQuant v2.4.14.0 interfaced with the Andromeda search engine against the UniProt human (v2024-26-08) database. Parameters for Max Quant were as follows: trypsin digestion, max missed cleavage site was set to two, precursor mass tolerance was set to 20 ppm, and fragment mass tolerance was set to 0.5 Daltons. The peptides were searched for the fixed modification of carbamidomethylation on cysteine, variable modifications of acetyl (Protein N-terminus), and the variable modifications of oxidation on methionine. The false discovery rate for both proteins and peptides was set to 0.01. Peptides were quantified based on unique and razor peptides with a label minimum ratio count of two. Label-free quantification was enabled with an LFQ min. ratio count of two. These filters are standard for proteomic label-free quantification analysis. Subsequent analysis was performed in Perseus v2.011.0. Principal component analysis plot and heatmap were generated in Metaboanalyst online software from log2 values based on normalized intensities **(Figure 7D and E)**.

## 6. Author contributions

Genome editing and X-chromosome screening was performed by J.W, Y.C., S.P., M.S., and M.N. NPCs were made by T.R. iPSCs were made at NYSCF with assistance from NYSCF Global Stem Cell Array® Team. The cardiomyocytes were derived by M.Y. and C.M., and the mass spectrometry was performed by R.B. and S.F. Project management administration at NYSCF was performed by C.H., C.M., L.B., F.J.M. and D.P. Overall project direction/supervision and funding acquisition was done by G.J.L. Blood and skin fibroblasts were collected and sent to NYSCF by E.M. and G.J.L. The initial manuscript was written by J.W. and team at NYSCF, followed by revision and addition of other data by R.M. and G.J.L.

## 7. Acknowledgements

We thank the families and the foundation, Ogden CARES for their participation and support. Maureen Gavin, Yessica Gonzalez, and Karen Amble assisted with family visits and/or blood collection at IBR. GJL thanks Jeff Talbot at Roseman University for facilitating the research on iPS-derived CMs in Utah. The RV-C2 antibody developed by Schiaffino S. et al^78^ was obtained from the Developmental Studies Hybridoma Bank (DSHB), created by the NICHD of the NIH and maintained at The University of Iowa, Department of Biology, Iowa City, IA 52242.

## NYSCF Global Stem Cell Array® Team^1^

Ankush Goyal

Anthony Chan

Barry McCarthy

Camille Fulmore

Christopher Hunter

Daniel White

Dong Woo Shin

Dillion Hutson

Farah Vejzagic

Geoff Buckley-Herd

Grayson Horn

Jenna Hall

John Cerrone

Jordan Goldberg

Kathryn Reggio

Katie Reggio

Kiran Ramnarine

Kola Campbell

Matt Green

Matthew Butawan

Matthew Zimmer

Michael Santos

Patrick Fenton

Paul McCoy

Peter Ferrarotto

Reid Otto

Ryan Kennedy

Saunil Dobariya

Sean DesMarteau

Selwyn Jacob

Siddharth Nimbalkar

Temi Oyelola

Lauren Bauer

Christopher Hunter

Connor McKnight

## 8. Ethical Approval

Both oral and written patient consent were obtained for creation of these cell lines, which are deidentified for distribution, with approval of protocol #7659 for the Jervis Clinic by the New York State Psychiatric Institute - Columbia University Department of Psychiatry Institutional Review Board.

## 10. Supplementary Information

**Table S1. List of iPSC lines available for generation**

**Table S2. Characterization and validation of iPSC lines**

**Table S3. Primers generated for sgRNA, PCR, ssODN, and their targets**

**Table S4. Overview of immunohistochemical staining protocol**

