## Supplementary Information for "A repository of Ogden syndrome patient derived iPSC lines and isogenic pairs by X-chromosome screening and genome-editing"

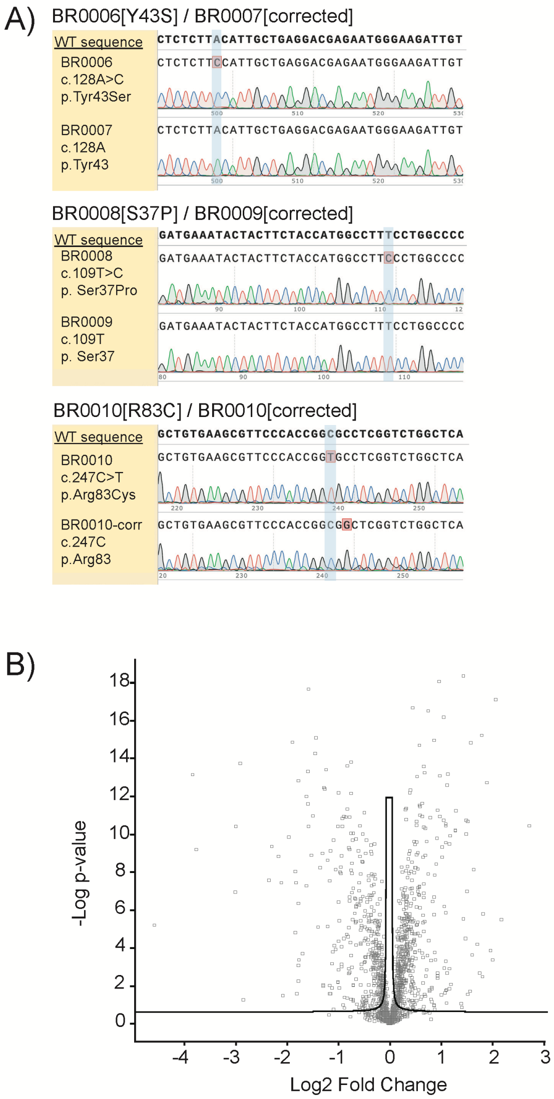


**Supplementary Figure 1. Validation of three male Ogden syndrome (OS) isogenic iPSC lines and characterization of one male OS isogenic pair.** Each mutation or correction was verified by amplifying the region of interest and sequenced using primers. See Figure 7. SnapGene was used to render the **A)** sequence chromatograms and alignment of wild-type NAA10 from the three isogenic Ogden syndrome lines. The mutations (red boxes) and corrections of interest are highlighted in blue. BR0010(R83corrected) was found to have a silent mutation that does not affect the primary protein sequence. **B)** BR0010(R83C) and BR0010(R83corrected) lines were further explored by differentiating into cardiomyocytes (CMs) and analyzed via LC-MS/MS. Volcano plot of quantified protein changes from the seven BR0010(R83C) with respect to BR0010(R83corrected). Each dot on the volcano plot represents an identified protein. The horizontal axis is the Log2 Fold Change (protein difference) between the BR0010(R83C) and BR0010(R83corrected). The vertical axis shows the -log p-value of the t-test. The curved line represents the FDR (false discovery rate) of 0.05 with an artificial group variance of 0.1.

**Supplementary Table 1. List of NAA10, NAA15, and related healthy control iPSC lines available for generation**

| NYSCF ID | Gene | Zygosity | Genotype / Family | Sex | Received Sample Type | Final Clone ID # to request cell line |
|---|---|---|---|---|---|---|
| BR0001* | *NAA10* | hemizygous | Ile72Thr, c.215 T>C | Male | PBMC | BR0001-01-MSV-419 |
| BR0002 | *NAA10* | heterozygous | Arg83Cys variant, WT on Xa. No R83C on Xa available | Female | PBMC | BR0002-01-MSV-031 |
| BR0003 | *NAA10* | heterozygous | Phe128Leu variant, F128L on Xa. No WT on Xa available | Female | PBMC | BR0003-01-MSV-066 |
| BR0004 | *NAA10* | heterozygous | Ala87Ser variant, WT on Xa. No A87S on Xa available | Female | PBMC | BR0004-02-MSV-083 |
| BR0005 | *NAA10* | heterozygous | Phe128Leu variant, F128L on Xa. No WT on Xa available. Mixed activation clones also observed and available. | Female | iPSC (NYSCF) | BR0005-01-MCS-168-EDRD0075 |
| BR0006 | *NAA10* | hemizygous | Tyr43Ser, older son, related to BR0030 and BR0031 | Male | iPSC (Stanford) | BR0006-04-MSV-021 |
| BR0007 | *NAA10* | hemizygous | Tyr43Ser Crispr-corrected to WT/y, from BR0006 | Male | iPSC (Stanford) | BR0007-04-MSV-075 |
| BR0008 | *NAA10* | hemizygous | Ser37Pro | Male | iPSC (Stanford) | BR0008-01-SV-001 |
| BR0009 | *NAA10* | hemizygous | Ser37Pro Crispr-corrected to WT/y from BR0008 | Male | iPSC (Stanford) | BR0009-01-SV-001 |
| BR0010 | *NAA10* | hemizygous | Arg83Cys | Male | PBMC | BR0010-01-MSV-163 |
| BR0010-EDIT | *NAA10* | hemizygous | Arg83Cys Crispr-corrected to WT/y, from BR0010 | Male | iPSC (NYSCF) | BR0010-01-MCS-421-EDIT0062B |
| BR0011 WT on Xa | *NAA10* | heterozygous | Arg83Cys variant, WT on Xa | Female | iPSC (NYSCF) | BR0011-01-MSV-081 |
| BR0011 R83C on Xa | *NAA10* | heterozygous | Arg83Cys variant, R83C on Xa | Female | iPSC (NYSCF) | BR0011-01-MSV-096 |
| BR0012 | *NAA10* | heterozygous | Leu121Val variant, WT on Xa. No L121V on Xa available. Mixed activation clones also observed and available. | Female | PBMC | BR0012-01-MSV-030 |
| BR0013 | *NAA10* | heterozygous | Arg83Cys, WT on Xa, no R83C on Xa available | Female | PBMC | BR0013-01-MSV-173 |
| BR0014 WT on Xa | *NAA10* | heterozygous | Phe128Leu variant, WT on Xa | Female | iPSC (NYSCF) | BR0014-01-MSV-046 |
| BR0014 F128L on Xa | *NAA10* | heterozygous | Phe128Leu variant, F128L on Xa | Female | iPSC (NYSCF) | BR0014-01-MSV-060 |
| BR0015 | *NAA10* | heterozygous | Arg83Cys variant, WT on Xa, No R83C on Xa available | Female | PBMC | BR0015-01-MSV-032 |
| BR0015-EDIT1 | *NAA10* | homozygous | BR0015 (Arg83Cys) Crispr-corrected to WT/WT from BR0015 | Female | iPSC (NYSCF) | BR0015-01-MCS-224-EDIT0138 |
| BR0015-EDIT 2 | *NAA10* | homozygous | BR0015 (Arg83Cys) Crispr-corrected to R83C/R83C from BR0015 | Female | iPSC (NYSCF) | BR0015-01-MCS-671-EDIT0138 |
| BR0016 WT on Xa | *NAA10* | heterozygous | Arg83Cys variant, WT on Xa | Female | iPSC (NYSCF) | BR0016-01-MCS-042-EDRD0057 |
| BR0016 R83C on Xa | *NAA10* | heterozygous | Arg83Cys variant, R83C on Xa | Female | iPSC (NYSCF) | BR0016-01-MCS-020-EDRD0057 |

| | | | | | | |
|---|---|---|---|---|---|---|
| BR0017 | *NAA10* | hemizygous | c.471+2T→A (splicing site of intron 7, removes 1 exon) | Male | Fibroblast | BR0017-01-MR-005 |
| BR0017-EDIT | *NAA10* | hemizygous | c.471+2T→A Crispr-corrected to WT/y | Male | iPSC (NYSCF) | BR0017-01-MCS-175-EDIT0118 |
| BR0018 | *NAA10* | hemizygous | c.471+2T→A (splicing site of intron 7, removes 1 exon), brother of BR0017 | Male | Fibroblast | BR0018-02-MR-031 |
| BR0019 | *NAA10* | hemizygous | c.471+2T→A (splicing site of intron 7, removes 1 exon), brother of BR0017 | Male | Fibroblast | BR0019-02-MR-012 |
| BR0030 | *NAA10* | | Tyr43Ser variant, from carrier mother for sons with BR0006 and BR0031. Unknown status for which allele on active X chromosome | Female | iPSC (Stanford) | BR0030-02-SV-001 |
| BR0031 | *NAA10* | hemizygous | Tyr43Ser, younger son, related to BR0030 | Male | iPSC (Stanford) | BR0031-02-SV-001 |
| BR0032 | *NAA10* | hemizygous | His034Tyr | Male | PBMC | BR0032-01-MSV-137 |
| BR0033 | *NAA15* | heterozygous | De novo NAA15 mutation. p.Asp0335Glufs*7 | Male | iPSC (Stanford) | BR0033-01-SV-001 |
| BR0034 | *NAA15* | heterozygous | Healthy sibling of BR0033-- without the variant, i.e. wild type | Male | iPSC (Stanford) | BR0034-01-SV-001 |

[*] BR is a random letter indicator at NYSCF for the repository of cells in the Matrix platform.
Grey shading: indicates isogenic male pairs of iPSCs, with currently 4 such pairs available.
Purple on grey shading: indicates isogenic female pairs of iPSCs, with one such pair available
Red lettering: female heterozygous pairs of clones where pathogenic variant is on Xa or WT is on Xa, where these lines are "isogenic", except Xa chromosome differs in each pair.

**Supplementary Table 2 Characterization and validation of iPSC lines**

| Classification | Test | Result | Data |
|---|---|---|---|
| Morphology | Celigo Bright Field | Characteristic iPSC morphology | Accessible through NYSCF |
| Phenotype | Quantitative analysis (gene expression analysis) | Positive for MYC, NANOG, SOX2, POU5F1 / Negative for Sox17, AFP, ANPEP, SRY | Accessible through NYSCF |
| | Qualitative analysis (immunocytochemistry) | N/A | |
| Genotype | Karyotype | Identical to parental | Accessible through NYSCF |
| Identity | Fluidigm / SNPTrace | Identical to parental line | Accessible through NYSCF |
| Mutation Analysis | Sanger sequencing | Confirmed correction | Accessible through NYSCF |
| Microbiology and virology | Mycoplasma results | Negative at 2 timepoints | Accessible through NYSCF |
| Differentiation potential | EB differentiation (Gene expression analysis) | Expression of 3 germ layer genes | Accessible through NYSCF |

**Supplementary Table 3 Primers generated for sgRNA, PCR, and ssODN and their targets**

| Name | Target | Forward/Reverse primer (5′-3′) |
|---|---|---|
| sgRNA | NAA10 R83C | tcagtttctgagccagaccg |
| PCR for sequencing: | NAA10 R83C | Fw cggttactgcaggggaacaa<br>Rw ttttcaggcagcagagggtg |
| ssODN | NAA10R83C | ccacctcttcctgacatgcagggagacatatttggcattgaagttctctatcatggctcgagaggcctggtccatcagtttctgagccagaccgaagcgccggtgggaacgcttcacagcctggtgg |
| sgRNA | NAA10 c.471+2T→A | cctcgtcggccatctgagtg |
| PCR for sequencing: | NAA10 c.471+2T→A | Fw caccttgttctcgatggcac<br>Rw acgataccaggagtggaggag |
| ssODN | NAA10 c.471+2T→A | tcgcctgcagccactgtcttggggctcctgagtgccgcccccgctcgccttgcttggcttcatgcaggcgcttAcctcgtcggccatctgagtgagAtcccgcttcatggcataggcgtcctcccca |

**Supplementary Table 4. Overview of immunohistochemical staining protocol**

| Reagent | Components | Concentration |
|---|---|---|
| 1. **Fixation Buffer** | 4% PFA | 1X |
| 1. **Permeabilization Buffer** | dPBS<br>TRITON-X-1001<br>HI-FBS | 1X<br>0.2%<br>1% |
| 1. **Wash Buffer** | dPBS<br>HI-FBS | 1X<br>1% |
| 1. **Blocking Buffer** | dPBS<br>BSA | 1X<br>5% |
| 1. **Live/Dead Stain** | Hoesht 33342 | 1mg/ml |
| 1. **Antibodies** | Anti-TRA-1-60-PE<br>Alexa fluro 488 Oct4 | 1:200<br>1:200 (0.5mg/ml) |
